# SNP2Cluster-Py: A pathogen-agnostic, context-aware framework for Core SNP and transmission-cluster analysis

**DOI:** 10.64898/2026.11.19.767809

**Authors:** Stanford Kwenda, Alan Christoffels

**Affiliations:** South African National Bioinformatics Institute, University of the Western Cape, Cape Town, South Africa

**Keywords:** Genomic epidemiology, Core SNP clustering, transmission-cluster detection, outbreak investigation, pathogen surveillance, research software

## Abstract

Whole-genome sequencing enables high-resolution identification of genetically related pathogen isolates, but genomic similarity alone does not establish epidemiologically meaningful transmission. We developed SNP2Cluster-Py, an open-source Python framework that separates context-aware Core SNP clustering from the subsequent application of epidemiological and temporal criteria. The software accepts a precomputed pairwise Core SNP-distance matrix and user-configured metadata and supports analysis of all eligible isolates in a single context, as well as sequence-type, geographical, epidemiological and ordered composite contexts. Within each context, authoritative Core clusters are defined as connected components of an SNP-threshold graph; K-means assignments and silhouette widths are retained as exploratory diagnostics only. We evaluated the workflow using 172 *Klebsiella pneumoniae* isolates from six neonatal-surveillance hospitals in South Africa, a focused 36-isolate single-hospital subset from the same surveillance dataset, and 168 *Streptococcus pneumoniae* carriage isolates from Ghana. In the full *K. pneumoniae* dataset, ST-only analysis assigned 141 isolates to Core SNP clusters, whereas Facility + ST analysis assigned 134, demonstrating how facility stratification can remove cross-facility genomic connections. In the single-hospital subset, both contexts assigned 30 of 36 isolates to five Core clusters, and Facility analysis retained 23 isolates in four temporally compatible clusters. In the pneumococcal dataset, ST-context analysis assigned 114 isolates to 38 Core clusters, which were characterised across sampling locations and time through Community analysis. SNP2Cluster-Py therefore provides a reproducible framework for adapting genomic clustering to specific surveillance questions while maintaining a clear distinction between genomic relatedness, analytically inferred transmission clusters and confirmed direct transmission. SNP2Cluster-Py v1.1.2 is available under the Apache License 2.0 through GitHub and Zenodo.

## Introduction

Whole-genome sequencing provides high-resolution information for identifying genetically related pathogen isolates and investigating possible transmission. In healthcare settings, genomic surveillance has detected clusters not identified through conventional infection-prevention surveillance and has distinguished transmission clusters from suspected outbreaks involving unrelated isolates [1–4]. However, genomic relatedness alone does not establish epidemiological linkage or direct transmission.

Interpretation of genomic similarity depends on the pathogen, analytical workflow, population structure, sampling density, surveillance setting, and period under investigation. SNP thresholds used to define related isolates vary across studies and organisms [1]. Healthcare genomic-surveillance studies frequently incorporate temporal and geographical linkage, but these criteria may not capture more complex transmission relationships [1,2]. Integrating genomic and epidemiological data provides a more informative basis for transmission-cluster detection. In a South African neonatal surveillance study, SNP-Epi clusters were defined as groups of genomically related isolates that also satisfied specified hospital and temporal-linkage criteria, based on pairwise SNP distances, hospital membership and specimen-collection dates [5]. Changing the SNP and temporal thresholds altered cluster detection, demonstrating the importance of explicit and configurable analytical criteria [5]. Prospective genomic-surveillance programmes have similarly shown that genomically detected clusters require epidemiological investigation before they can guide infection-prevention interventions [2–4].

Analytical tools are therefore needed that reproducibly integrate genomic distances with epidemiological metadata while retaining the context in which clusters were identified. This is particularly relevant in resource-limited settings, where sustainable pathogen-genomics implementation depends on accessible analytical workflows addressing locally relevant public-health questions [6].

Existing software spans upstream variant calling, integrated genomic-surveillance platforms, phylogenetic outbreak reconstruction, dynamic surveillance-cluster detection and interactive visualization [9–13]. These tools address overlapping but distinct stages of bacterial genomic epidemiology, ranging from sequence processing and phylogenetic analysis to longitudinal cluster detection and communication of genomic findings.

SNP2Cluster-Py is an open-source Python reimplementation and extension of the original SNP2Cluster method [7]. The framework performs Core SNP clustering from a precomputed pairwise SNP-distance matrix and can subsequently integrate Core cluster membership with epidemiological grouping and specimen-collection dates. Enhancements include configuration-driven execution, flexible metadata mapping, ordered Core clustering contexts, Facility, Area and Community analysis modes, automated testing, machine-readable outputs, run provenance, interactive visualizations and scientific HTML reporting.

A central feature of SNP2Cluster-Py is the separation of the context defining which isolates are eligible to undergo Core SNP clustering together from the epidemiological and temporal criteria used in subsequent transmission analysis. Within each context, the configured SNP cutoff defines an undirected threshold graph, and connected components containing at least two isolates define authoritative Core clusters. This allows genomic clusters to be defined globally, within sequence types, within epidemiological or geographical groups, or within ordered combinations of these variables.

Here we describe SNP2Cluster-Py and demonstrate its context-aware analytical workflow using two previously published genomic-surveillance datasets: 172 *Klebsiella pneumoniae* isolates collected across six hospitals through South African neonatal invasive-infection surveillance [5], including a focused 36-isolate single-hospital subset, and 168 *Streptococcus pneumoniae* carriage isolates collected across community sampling locations in Ghana [8]. The full *K. pneumoniae* dataset was used to compare threshold-graph Core SNP clustering under ST-only and Facility + ST contexts, while the single-hospital subset provided a controlled comparison in which facility was constant and supported the subsequent Facility-level transmission analysis. The pneumococcal dataset was used to demonstrate ST-context Core clustering followed by Community analysis across sampling locations and time.

## Implementation

### Software design

SNP2Cluster-Py v1.1.2 was developed in Python 3.10 or later as an open-source command-line framework for context-aware Core SNP clustering and epidemiologically informed downstream cluster analysis. The software is a Python reimplementation and extension of the original SNP2Cluster method [7]. Analyses are executed through the snp2cluster command-line interface and controlled using YAML configuration files.

SNP2Cluster-Py uses pandas and NumPy for data processing, SciPy and scikit-learn for exploratory clustering diagnostics, NetworkX and PyVis for graph-based analysis and interactive visualization, Matplotlib and seaborn for static visualization, and Typer and Rich for the command-line interface. The software is distributed under the Apache License 2.0.

### Inputs and configuration

The principal input is a square pairwise Core SNP-distance matrix with matching row and column sample identifiers. Depending on the analysis, users may additionally provide epidemiological metadata, specimen-collection dates and multilocus sequence-typing profiles or other lineage assignments. Supported tabular formats include CSV, TSV, Excel and Parquet.

The YAML configuration specifies the input files, sample and metadata fields, analysis mode, ordered Core-clustering context, SNP and temporal thresholds, bootstrap count, random seed, date format and requested outputs. Input validation checks and normalizes the supplied data, reconciles sample identifiers across files, verifies the structure and symmetry of the SNP-distance matrix, and confirms the availability of variables required for the selected analysis.

### Analytical workflow

SNP2Cluster-Py separates Core SNP clustering from downstream Facility, Area or Community analysis. Eligible isolates are first partitioned into user-defined ordered Core contexts. Within each context, the configured SNP cutoff defines graph edges, and connected components determine authoritative Core-cluster membership. Downstream analysis is optional and subsequently combines Core-cluster membership with configured epidemiological groupings and specimen-collection dates. This design distinguishes genomic relatedness within a specified population from clusters that additionally satisfy epidemiological and temporal criteria.

### Context-aware Core SNP clustering

The ordered core_cluster_context parameter determines which isolates are eligible to undergo Core SNP clustering together. Supported contexts include all eligible isolates in a single context, sequence type, a primary metadata variable, a secondary metadata variable and ordered combinations of these variables. Isolates assigned to different contexts are analysed independently, and the configured context order is retained in the resulting canonical cluster identifiers.

Within each context, isolates are represented as nodes in an undirected SNP-threshold graph. An edge is created between two isolates when their pairwise Core SNP distance is less than or equal to the configured cutoff. Connected components containing at least two isolates are retained as authoritative Core SNP clusters.

This connected-component procedure implements single-linkage clustering. Isolates can therefore belong to the same Core cluster through a chain of qualifying pairwise links even when the direct distance between some non-adjacent members exceeds the configured cutoff. Candidate values of *k*, K-means assignments and silhouette widths are generated as exploratory diagnostics and do not split, merge, retain or exclude authoritative Core-cluster members. Each retained cluster receives a canonical identifier combining its ordered Core context with a context-specific cluster number.

### Transmission-cluster analysis

Transmission analysis applies epidemiological and temporal criteria to isolates assigned to Core SNP clusters. Facility mode evaluates clustering within individual facilities or equivalent local units, whereas Area mode evaluates clustering within broader geographical or organizational groupings. In both modes, isolates remain constrained by the lineage and other ordered contexts established during Core SNP clustering.

Community mode evaluates the temporal and geographical distribution of a specified lineage across multiple sampling locations. In carriage or population-surveillance datasets, Community assignments may be interpreted as putative strain-sharing clusters rather than confirmed transmission events.

Downstream cluster membership indicates that isolates are connected through the configured genomic, epidemiological and temporal criteria. It does not establish direct transmission, transmission direction, source attribution or the absence of unsampled intermediates.

### Outputs and reproducibility

SNP2Cluster-Py generates machine-readable Core and downstream cluster assignments, complete isolate-level annotation tables, exploratory K-means and silhouette diagnostics, annotated Core SNP heatmaps, temporal scatterplots, interactive Core minimum spanning trees, downstream cluster networks, validation and audit records, a run manifest and a self-contained scientific HTML report.

The run manifest and HTML report record the input files, effective analytical parameters, configured columns, validation status, software version, operating environment and dependency versions. Core minimum spanning trees summarize genetic relationships among eligible isolates. Downstream network edges represent within-cluster minimum-spanning-tree relationships labelled by SNP distance; edge styling can additionally indicate whether the direct collection-date difference is within the configured temporal cutoff. These edges support visualization of analytically defined clusters and must not be interpreted as evidence of confirmed direct or directional transmission.

## Example use cases

### South African neonatal Klebsiella pneumoniae surveillance

The first use case comprised 172 *K. pneumoniae* isolates collected across six hospitals through the Baby GERMS-SA neonatal invasive-infection surveillance programme in South Africa [5]. Pairwise Core SNP distances were analysed together with hospital, ward-type, sequence-type and specimen-collection-date metadata. Following the thresholds used in the source study, Core genomic relatedness was assessed using a maximum distance of 11 SNPs, and Facility-level transmission clusters were identified using a 14-day temporal window.

The complete dataset was analysed under two Core-clustering contexts. The first analysis grouped isolates by sequence type using core_cluster_context: [ST], allowing isolates of the same ST from different hospitals to contribute to the same SNP-threshold connected component. The second analysis used core_cluster_context: [MainVar, ST], with hospital configured as the primary variable, thereby restricting Core clustering to isolates sharing both hospital and ST. Facility-level transmission analysis was subsequently applied to the Facility + ST Core assignments. These analyses were used to assess how facility stratification affected Core-cluster membership across the six-hospital dataset.

A focused subset of 36 isolates from one hospital was analysed separately under [ST] and [MainVar, ST] contexts using the same SNP-distance matrix and metadata. Because hospital was constant within this subset, the comparison tested whether adding the facility variable changed the underlying Core-cluster membership or only the canonical context labels. The subset was also used to demonstrate how the 14-day temporal criterion refined genomic Core clusters into Facility-level transmission clusters.

### Ghanaian community pneumococcal carriage surveillance

The second use case comprised 168 *Streptococcus pneumoniae* carriage isolates from a published genomic study conducted across community sampling locations in Cape Coast, Ghana [8]. Pairwise Core SNP distances were analysed together with sequence-type, sampling-location and collection-date metadata. Core SNP clustering was performed within STs using core_cluster_context: [ST] and a maximum pairwise-link threshold of 12 SNPs.

Community analysis subsequently evaluated the temporal and geographical distribution of the ST-specific Core-clustered isolates across sampling locations using a 45-day temporal window. Because the dataset represented pneumococcal carriage surveillance rather than a defined outbreak investigation, retained groups were interpreted as putative strain-sharing clusters. These assignments indicate that isolates were connected through the configured genomic, lineage, geographical and temporal criteria, but do not establish direct host-to-host transmission, directionality or source attribution.

## Results

### SNP2Cluster-Py produced context-aware analytical and reporting outputs

SNP2Cluster-Py v1.1.2 completed ST-only and Facility + ST Core analyses of the full 172-isolate *Klebsiella pneumoniae* dataset, focused validation analyses of the 36-isolate single-hospital subset, and ST-context Core and Community analyses of the 168-isolate *Streptococcus pneumoniae* carriage dataset. Each analysis produced machine-readable cluster assignments, exploratory K-means and silhouette diagnostics, annotated Core SNP heatmaps, interactive minimum spanning trees and a scientific HTML report. Transmission-mode analyses additionally produced temporal scatterplots, downstream cluster assignments, transmission networks and the corresponding machine-readable node and edge tables.

### Facility stratification reduced Core-cluster membership in the full K. pneumoniae dataset

Under ST-only Core clustering, 141 of 172 *K. pneumoniae* isolates (82.0%) were assigned to Core SNP clusters. Adding facility to the ordered Core context reduced Core-cluster membership to 134 isolates (77.9%). Isolate-level comparison showed that all 134 isolates clustered under the Facility + ST context were also clustered under the ST-only context. Seven isolates that clustered under ST-only analysis became unclustered after facility stratification, no isolates became newly clustered, and 31 isolates remained unclustered under both configurations (Figure 2).

**Figure 1.**
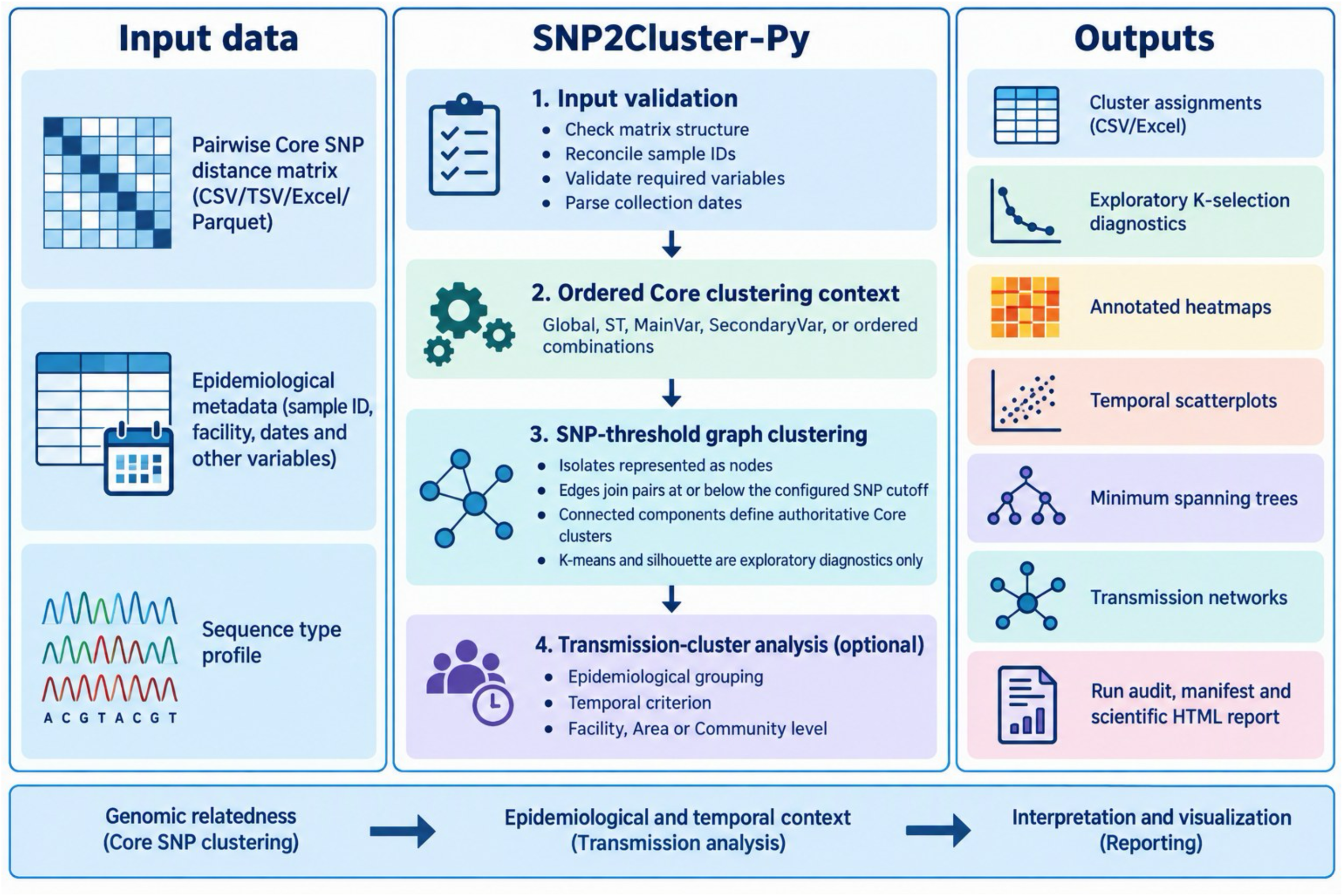
SNP2Cluster-Py analytical workflow. A pairwise Core SNP-distance matrix is combined with epidemiological metadata and, where applicable, sequence-type information. Following input validation, isolates are partitioned according to the specified ordered Core clustering context. Within each context, isolate pairs at or below the configured SNP cutoff are linked in an undirected graph, and connected components containing at least two isolates define the authoritative Core SNP clusters. K-means assignments and silhouette widths are retained as exploratory diagnostics only. Optional Facility, Area or Community analysis subsequently applies epidemiological and temporal criteria. Outputs include cluster assignments, exploratory diagnostics, annotated heatmaps, temporal scatterplots, minimum spanning trees, transmission networks, run-provenance records and a scientific HTML report.

**Figure 2.**
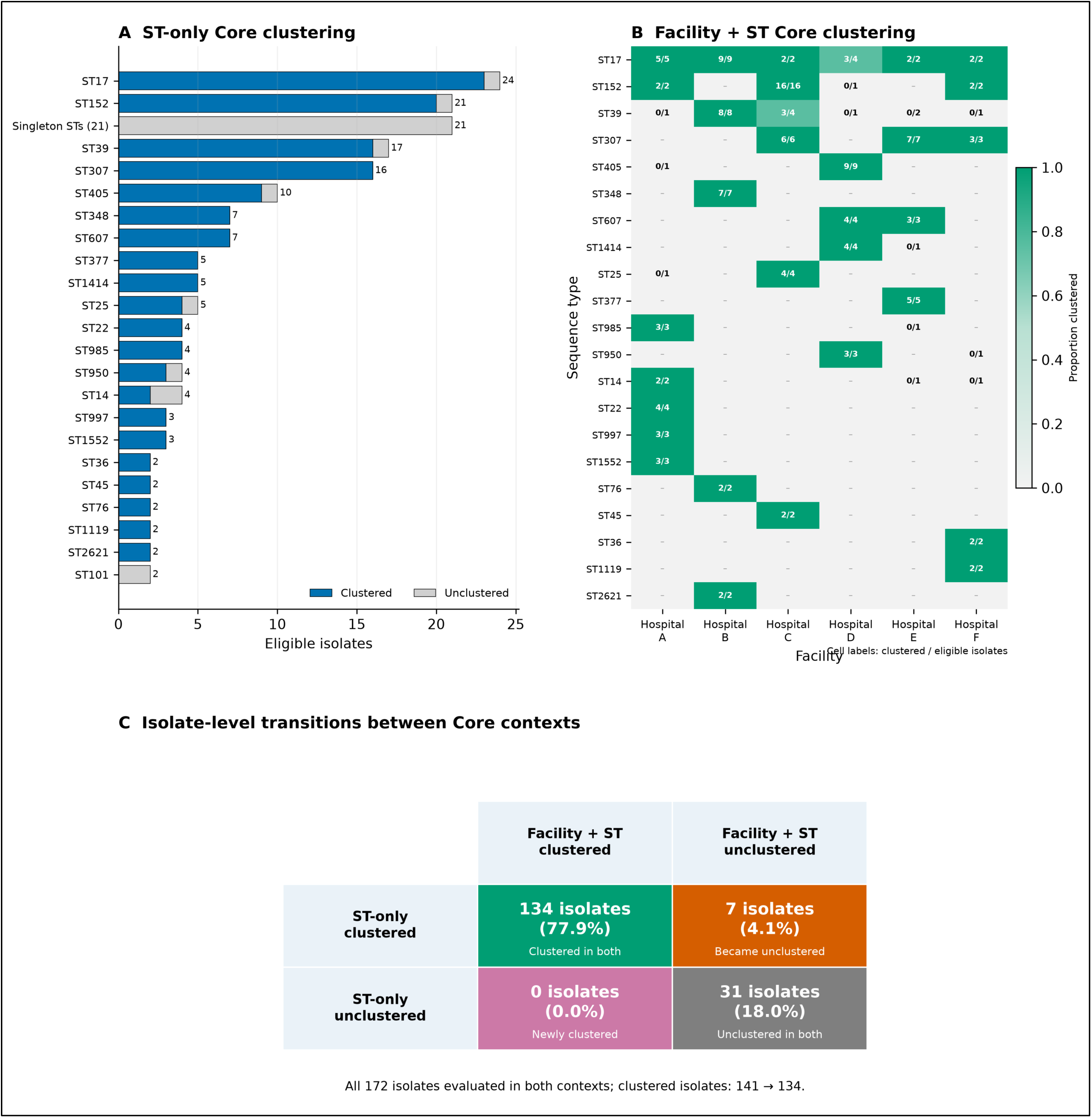
Effect of Core clustering context on Klebsiella pneumoniae cluster membership. **(A)** ST-only Core clustering among 172 isolates. Horizontal bars show the numbers of Core-clustered and unclustered isolates within the most frequently represented sequence types; sequence types represented by a single isolate are combined as singleton STs. **(B)** Facility + ST Core clustering. Cells show the number of Core-clustered isolates relative to the number of eligible isolates within each facility–ST context, with colour indicating the proportion clustered. **(C)** Isolate-level transitions between the two Core contexts. Of 172 isolates evaluated under both configurations, 134 remained clustered, seven were clustered under ST-only analysis but became unclustered after facility stratification, no isolates became newly clustered, and 31 remained unclustered under both configurations. Core membership was defined using authoritative SNP-threshold connected components.

This reduction reflected the smaller eligible isolate sets created by the Facility + ST context. Under ST-only analysis, SNP-threshold links could connect isolates of the same sequence type across different facilities. Under Facility + ST analysis, isolates could form Core clusters only through qualifying links to isolates sharing both sequence type and facility. Facility stratification therefore separated seven isolates whose ST-only Core-cluster membership depended on cross-facility genomic connections.

### ST and Facility + ST contexts produced equivalent Core membership in the single-hospital subset

The 36-isolate single-hospital subset provided a controlled comparison in which facility was constant. The ST-only analysis partitioned eligible isolates by sequence type, whereas the Facility + ST analysis partitioned isolates by the ordered combination of hospital and sequence type. Because all isolates originated from the same hospital, both configurations created the same eight ST-defined Core contexts and produced identical Core-cluster membership. The configurations differed only in their canonical identifiers: ST-only contexts used labels such as 307__1, whereas Facility + ST contexts used labels such as Hospital C__307__1.

Both configurations assigned 30 of 36 isolates (83.3%) to five Core SNP clusters corresponding to ST17, ST25, ST45, ST152 and ST307. Six isolates remained outside retained Core clusters, and cluster sizes ranged from two to 16 isolates. All six ST307 isolates formed one SNP-threshold connected component at the 11-SNP cutoff under both configurations. These findings confirmed that adding a constant facility variable changed the canonical context labels but not the underlying SNP-threshold connected components.

### Facility analysis refined *K. pneumoniae* Core clusters using temporal criteria

Facility-level analysis applied the 14-day temporal criterion to isolates assigned to Facility + ST Core clusters defined at the 11-SNP cutoff. Temporal scatterplots retained Core-clustered isolates that did not satisfy the temporal criterion as epidemiological context. Four Facility transmission clusters were identified, containing 23 of the 30 Core-clustered isolates: 15 ST152 isolates, four ST307 isolates, two ST17 isolates and two ST45 isolates. Seven Core-clustered isolates remained transmission-unclustered because they were not connected through the configured 14-day temporal criterion.

The ST307 context illustrates the distinction between genomic and transmission clustering. All six ST307 isolates formed one Core SNP cluster because they belonged to the same connected component at the 11-SNP cutoff. Four isolates were connected through consecutive temporal intervals of 7, 1 and 5 days and formed one Facility transmission cluster. The two remaining ST307 isolates remained members of the Core cluster but were excluded from the transmission cluster because they were separated from the central temporal chain by intervals of 48 and 42 days. The resulting network summarizes within-cluster genetic relationships and does not establish direct transmission, transmission direction or source attribution (Figure 3).

**Figure 3.**
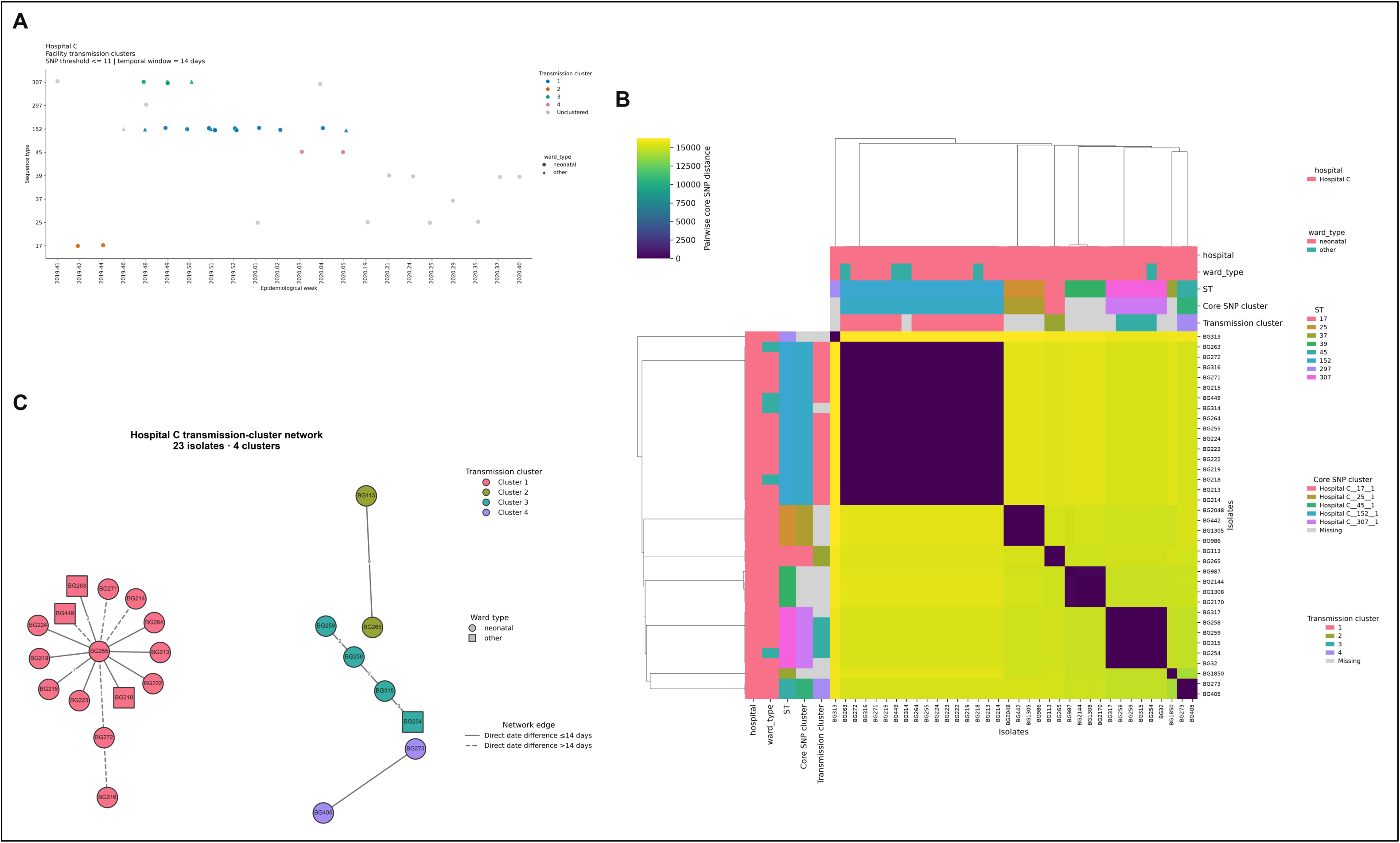
Facility-level analysis of *Klebsiella pneumoniae* isolates from the single-hospital subset. **(A)** Temporal distribution of four Facility transmission clusters and transmission-unclustered isolates across the represented sequence types. Core-clustered isolates that did not satisfy the temporal criterion are retained to provide epidemiological context. **(B)** Pairwise Core SNP-distance heatmap showing genomic relationships and Facility + ST Core-cluster assignments among 36 isolates. Thirty isolates were assigned to five Core SNP clusters. **(C)** Transmission-cluster network showing 23 isolates in four clusters identified using an 11-SNP threshold and a 14-day temporal window. Node shapes indicate ward type, node colours indicate transmission-cluster assignment, and edges show within-cluster minimum-spanning-tree relationships labelled by SNP distance. Solid edges connect isolates whose direct collection-date difference is within 14 days, whereas dashed edges connect isolates whose direct date difference exceeds 14 days. Network edges support visual interpretation and do not imply direct or directional transmission.

### Community mode identified pneumococcal strain-sharing patterns across sampling locations

Community analysis applied a 12-SNP threshold and a 45-day temporal window to isolates assigned to ST-specific Core SNP clusters. Of the 168 pneumococcal isolates analysed, 114 were assigned to 38 Core SNP clusters and retained in 38 Community strain-sharing clusters. Each represented ST contributed one retained Community cluster; no ST contributed multiple Community clusters under the configured criteria.

ST802 Community cluster 1 comprised 12 isolates collected at six locations: AME, DAN, GS, H1, NEW and UIC. The clustered isolates were distributed between epidemiological weeks 2018.04 and 2018.08, indicating the presence of a genomically related lineage across multiple sampling locations and collection times. One additional ST802 isolate from UIC remained outside the Community cluster despite belonging to the same ST.

ST5103 Community cluster 1 comprised seven isolates collected at five locations: GS, IMA, KAK, NEW and PEDU. These isolates were retained as one Community cluster under the 12-SNP and 45-day criteria, providing a second example of a genomically related carriage lineage distributed across several sampling locations (Figure 4).

**Figure 4.**
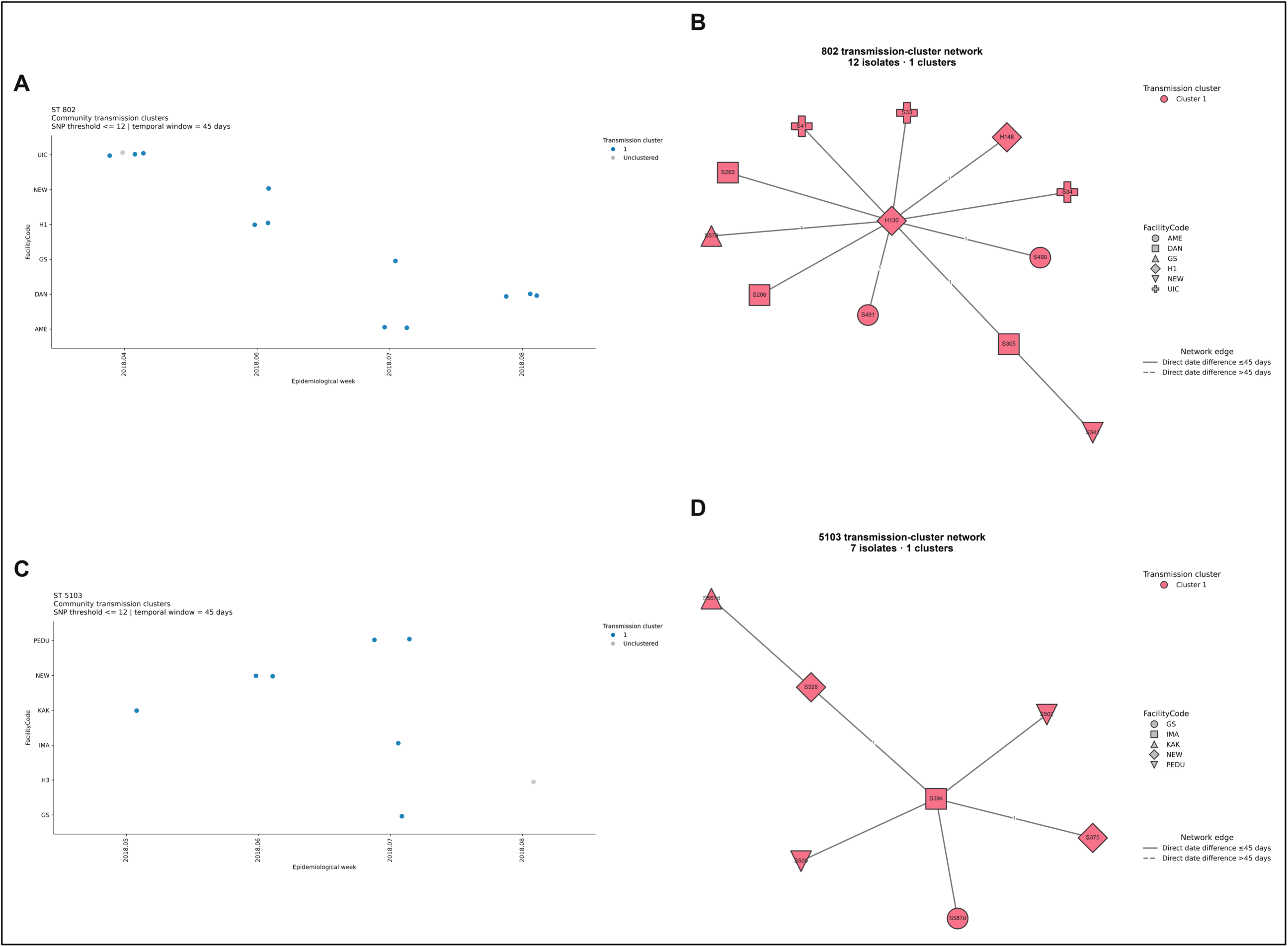
Community-level analysis of pneumococcal carriage isolates across sampling locations. **(A)** Temporal and geographical distribution of ST802 isolates across AME, DAN, GS, H1, NEW and UIC. **(B)** Genomic relationships among 12 transmission-clustered ST802 isolates represented as a single Community network spanning six sampling locations. **(C)** Temporal and geographical distribution of ST5103 isolates across GS, IMA, KAK, NEW and PEDU. **(D)** Genomic relationships among seven transmission-clustered ST5103 isolates represented as a single Community network spanning five sampling locations. Community clusters were identified using a 12-SNP threshold and a 45-day temporal window. Node colours indicate Community-cluster assignment, node shapes indicate sampling location, and edges show within-cluster minimum-spanning-tree relationships labelled by SNP distance. Solid edges connect isolates whose direct collection-date difference is within 45 days, whereas dashed edges connect isolates whose direct date difference exceeds 45 days. Network edges support visual interpretation and do not imply direct or directional transmission.

Together, the ST802 and ST5103 examples demonstrate how Community analysis can identify ST-specific genomic clusters distributed across locations and time. Because the dataset represented pneumococcal carriage surveillance and did not include individual-level contact, movement or exposure information, the identified groups were interpreted as putative strain-sharing clusters rather than confirmed direct-transmission events. Community-cluster membership does not establish transmission direction, source attribution or the absence of unsampled intermediates.

### Summary of the genomic clustering and downstream analyses

Across both applications, the Core-clustering context determined which isolates were eligible to cluster together, while Facility and Community analyses applied the configured epidemiological and temporal criteria to isolates assigned to Core SNP clusters. In the full 172-isolate *Klebsiella pneumoniae* dataset, ST-only analysis assigned 141 isolates (82.0%) to 25 Core SNP clusters across 21 retained ST contexts. Facility + ST analysis assigned 134 isolates (77.9%) to 32 Core SNP clusters across 32 retained facility–ST contexts. Subsequent Facility analysis retained 84 isolates in 24 temporally compatible clusters across the six hospitals.

In the 36-isolate single-hospital subset, ST-only and Facility + ST contexts created the same eight eligible contexts and five Core SNP clusters containing 30 isolates. Subsequent Facility analysis retained 23 of the 30 Core-clustered isolates in four temporally compatible clusters. The equivalent Core membership under ST-only and Facility + ST analysis confirmed that adding a constant facility variable changed the canonical context labels but not the underlying SNP-threshold connected components.

In the pneumococcal carriage dataset, ST-context Core analysis assigned 114 of 168 isolates (67.9%) to 38 Core SNP clusters across 38 retained ST contexts. All 114 Core-clustered isolates were retained in 38 Community strain-sharing clusters under the configured 12-SNP and 45-day criteria. Summary statistics for the six analyses are presented in Table 1.

**Table 1.** Summary of the SNP2Cluster-Py use-case analyses.

| Dataset | Core context and downstream analysis | Eligible isolates | Core contexts, eligible/retained | Core SNP clusters | Core-clustered isolates, n (% of eligible) | Core cluster size, range | Downstream clusters | Downstream-clustered isolates, n (% of eligible) | Downstream cluster size, range |
|---|---|---|---|---|---|---|---|---|---|
| Full *K. pneumoniae* dataset | ST-only Core analysis | 172 | 43/21 | 25 | 141 (82.0%) | 2–23 | — | — | — |
| Full *K. pneumoniae* dataset | Facility + ST Core analysis; Facility analysis | 172 | 67/32 | 32 | 134 (77.9%) | 2–16 | 24 | 84 (48.8%) | 2–15 |
| Single-hospital *K. pneumoniae* subset | ST-only Core analysis | 36 | 8/5 | 5 | 30 (83.3%) | 2–16 | — | — | — |
| Single-hospital *K. pneumoniae* subset | Facility + ST Core analysis; Facility analysis | 36 | 8/5 | 5 | 30 (83.3%) | 2–16 | 4 | 23 (63.9%) | 2–15 |
| *S. pneumoniae* carriage dataset | ST-only Core analysis | 168 | 67/38 | 38 | 114 (67.9%) | 2–12 | — | — | — |
| *S. pneumoniae* carriage dataset | ST-only Core analysis; Community analysis | 168 | 67/38 | 38 | 114 (67.9%) | 2–12 | 38 | 114 (67.9%) | 2–12 |

ST, sequence type. Eligible Core contexts include all context groups evaluated during Core clustering; retained Core contexts contain at least one authoritative Core SNP cluster. Downstream clusters refer to Facility transmission clusters for K. pneumoniae and Community strain-sharing clusters for S. pneumoniae. Percentages in the table are calculated relative to all eligible isolates in the corresponding analysis. In the full Facility + ST analysis, 84 isolates represented 48.8% of all 172 eligible isolates and 62.7% of the 134 Core-clustered isolates. In the single-hospital Facility analysis, 23 isolates represented 63.9% of all 36 eligible isolates and 76.7% of the 30 Core-clustered isolates. Facility analysis used an 11-SNP threshold and a 14-day temporal window; Community analysis used a 12-SNP threshold and a 45-day temporal window.

## Discussion

SNP2Cluster-Py provides a configuration-driven framework that separates the isolates eligible for joint Core SNP clustering from the subsequent application of epidemiological and temporal criteria. This distinction allows the same genomic dataset to address different surveillance questions depending on whether all eligible isolates are considered together or divided by lineage, facility, geographical grouping or an ordered combination of these variables. Within each defined context, authoritative Core SNP clusters are identified as connected components of an SNP-threshold graph, while epidemiological and temporal criteria are applied only during the subsequent Facility, Area or Community analysis.

The complete *K. pneumoniae* dataset demonstrated how the choice of Core-clustering context can alter both cluster membership and cluster structure. Under ST-only analysis, 141 of 172 isolates were assigned to 25 Core SNP clusters. Adding facility to the context reduced membership to 134 isolates but increased the number of separately labelled Core clusters to 32. This increase in cluster count did not represent increased clustering. Instead, facility stratification divided ST-defined populations into smaller hospital–ST contexts, preventing cross-facility SNP-threshold links and separating components that had been connected when isolates of the same ST were analysed together across hospitals. Seven isolates consequently lost Core-cluster membership, while no previously unclustered isolate became clustered.

The single-hospital subset provided a controlled comparison of the same two contexts. Because facility was constant, ST-only and Facility + ST analyses created the same eight eligible isolate groups and produced identical membership: 30 of 36 isolates were assigned to five Core SNP clusters. Only the canonical identifiers changed to preserve the explicitly configured context. This result confirms that adding a context variable does not inherently alter cluster membership. Membership changes only when the variable partitions the eligible isolates into analytically distinct contexts.

Facility analysis then applied the temporal criterion to the Facility + ST Core assignments. Across the complete six-hospital dataset, 84 of 134 Core-clustered isolates were retained in 24 Facility transmission clusters. In the focused single-hospital subset, 23 of 30 Core-clustered isolates were retained in four temporally compatible clusters. The ST307 example illustrates the distinction between genomic clustering and temporally informed transmission clustering: all six ST307 isolates formed one Core SNP cluster, but only four were connected through the configured 14-day temporal criterion. Core SNP clustering therefore identifies genomic relatedness within the selected context, whereas Facility analysis identifies the subset that also satisfies the configured epidemiological and temporal conditions.

The pneumococcal carriage analysis demonstrated a different downstream application. ST-context Core analysis assigned 114 of 168 isolates to 38 Core SNP clusters, and all 114 were retained in 38 Community strain-sharing clusters under the 12-SNP and 45-day criteria. The ST802 and ST5103 examples showed that genomically related carriage lineages could span multiple sampling locations and epidemiological weeks. Unlike the *K. pneumoniae* Facility analysis, Community analysis did not reduce membership among the Core-clustered isolates in this dataset. This difference reflects the interaction between the selected Core context, the distribution of isolates across sampling locations and time, and the configured analytical thresholds. Because the pneumococcal dataset represented carriage surveillance and lacked individual-level contact, movement and exposure information, the retained groups were interpreted as putative strain-sharing clusters rather than confirmed transmission events.

### Positioning alongside related software

Several existing tools support complementary stages of bacterial genomic epidemiology. For example, Snippy performs reference-based variant calling and Core SNP alignment and generates upstream genomic data from which pairwise SNP distances are derived [9]. P-DOR provides an end-to-end outbreak-reconstruction workflow that selects related public background genomes, constructs an SNP-based phylogeny, identifies threshold-defined clusters and integrates patient metadata into spatiotemporal outputs [10]. DODGE was designed for continuously expanding surveillance datasets and dynamically selects cluster-specific genetic thresholds using temporal criteria and the preceding genomic background [11]. CATHAI supports interactive exploration of genomic-distance relationships alongside clinical and epidemiological metadata [12], while Pathogenwatch provides a broader genomic-surveillance platform incorporating genome analysis, lineage assignment, antimicrobial-resistance and virulence characterization, global contextualization and interactive visualization [13].

SNP2Cluster-Py occupies a narrower downstream analytical position. It does not perform read processing, genome assembly, variant calling, recombination filtering or phylogenetic inference. Instead, it accepts a precomputed pairwise Core SNP-distance matrix and provides explicit control over the biological and epidemiological contexts within which isolates are eligible to cluster. Its principal contribution is the separation of ordered, context-aware Core SNP clustering from the subsequent application of epidemiological and temporal criteria. It also produces machine-readable cluster assignments, complete isolate-level annotation outputs, interactive visualizations, scientific HTML reports and structured run-provenance records. SNP2Cluster-Py is therefore intended to complement, rather than replace, upstream genomic-analysis and broader surveillance platforms.

### Strengths and limitations

The principal strengths of SNP2Cluster-Py are its explicit ordered-context model, authoritative SNP-threshold graph definition, separation of Core and downstream analysis, configuration-controlled execution, machine-readable outputs and structured run provenance. The framework retains eligible unclustered isolates in its complete annotation outputs, allowing retained clusters to be interpreted against the wider genomic and epidemiological background. Exploratory K-means assignments and silhouette widths remain available for examining the structure of SNP-distance profiles without controlling authoritative Core-cluster membership. The same analytical architecture can be applied at Facility, Area or Community level by changing the configured context and downstream grouping variables.

The analysis nevertheless depends on the quality, comparability and representativeness of the precomputed SNP-distance matrix and accompanying metadata. Results may be affected by reference selection, variant-calling and recombination-filtering procedures applied before the distance matrix is generated. Cluster assignments are also sensitive to the selected context variables, SNP thresholds and temporal intervals. These parameters should therefore be justified for the organism, population and surveillance question rather than treated as universally transferable.

The connected-component definition implements single-linkage clustering. Isolates may consequently belong to the same Core cluster through a chain of qualifying pairwise links even when the direct distance between some non-adjacent members exceeds the configured SNP threshold. Cluster diameter, internal distance distributions and network structure should therefore be inspected when interpreting large or heterogeneous components. In addition, Facility, Area and Community assignments identify isolates that satisfy configured genomic, epidemiological and temporal criteria but cannot establish direct transmission, transmission direction, source attribution or the absence of unsampled intermediates.

The present study demonstrates the workflow using two previously published bacterial surveillance datasets but does not provide formal comparative benchmarking against alternative clustering frameworks. It also does not evaluate sensitivity across alternative SNP thresholds, temporal windows, reference genomes or upstream variant-calling pipelines. Future work should compare SNP2Cluster-Py with alternative genomic-clustering approaches, quantify the stability of assignments across analytical parameters and assess performance in prospectively expanding surveillance datasets.

## Conclusions

SNP2Cluster-Py v1.1.2 provides an open-source, context-aware framework for Core SNP clustering and epidemiologically informed downstream analysis. Ordered Core contexts define which isolates are eligible to cluster together, SNP-threshold connected components determine authoritative Core-cluster membership, and optional Facility, Area and Community modes apply epidemiological and temporal criteria.

In the complete *Klebsiella pneumoniae* dataset, facility stratification reduced Core-cluster membership by preventing cross-facility genomic connections, whereas the controlled single-hospital comparison showed that adding a constant facility variable changed canonical identifiers without altering membership. Facility analysis subsequently identified temporally compatible subsets of the Core-clustered isolates. In the pneumococcal carriage dataset, Community analysis characterised ST-specific genomic clusters distributed across sampling locations and time.

SNP2Cluster-Py therefore enables genomic clustering to be adapted reproducibly to different surveillance populations and questions while maintaining clear distinctions between genomic relatedness, analytically inferred transmission or strain-sharing clusters and confirmed direct transmission.

## Supporting information

Supplementary HTML Files

## Ethics statement

The use cases were secondary analyses of previously published, de-identified datasets. Ethics approvals and consent procedures for the original studies are described in the corresponding publications [5,8]. No additional identifiable participant information was accessed or analysed.

## Data availability

The use-case datasets were derived from previously published studies [5,8]. Public sequence data for the pneumococcal carriage study are available under NCBI BioProject PRJNA1100994. Access conditions for the Baby GERMS-SA genomic data are described in the corresponding publication [5].

Derived analytical inputs permitted for redistribution, de-identified metadata, YAML configuration files, machine-readable output tables, static figures, interactive visualizations and scientific HTML reports are available through the SNP2Cluster-Py manuscript reproducibility repository. Materials that cannot be redistributed are documented with their access conditions, data dictionaries and instructions for reconstructing the corresponding analyses. A versioned archive of the final reproducibility materials will be deposited in Zenodo, and the DOI will be added before submission.

## Code availability

SNP2Cluster-Py v1.1.2 is available from GitHub under the Apache License 2.0. The software record is archived in Zenodo under DOI 10.5281/zenodo.23045861. The original R-based SNP2Cluster implementation is archived separately under DOI 10.5281/zenodo.14060296.

The YAML configurations, figure-generation scripts, run instructions, and other materials required to reproduce the manuscript analyses are available from the SNP2Cluster-Py bioRxiv reproducibility repository. A versioned Zenodo archive of the final reproducibility repository will be created before submission, and its DOI will be added to the manuscript.

## Acknowledgements

The authors acknowledge the investigators, laboratory teams, participating institutions and study participants responsible for generating and curating the Baby GERMS-SA and Ghana pneumococcal carriage datasets used in the example analyses.

Microsoft Copilot was used to assist with manuscript language editing, preparation of the analytical workflow diagram, and development of Python scripts used to generate or compose Figures 2–4 and selected supplementary materials. The authors reviewed and verified all manuscript text, analytical values, code, visualizations and interpretations and take full responsibility for the final work.

## Author contributions

Conceptualization: SK. Methodology, software and validation: SK. Writing, original draft: SK. Writing, review and editing: SK and AC. Supervision: AC.

## Supplementary information

The supplementary information comprises five static supplementary figures, an Excel workbook containing Tables S1–S5 and a ZIP archive containing interactive HTML visualizations and their associated machine-readable tables. Complete reproducibility materials are provided through the companion repository and its versioned archive. The reproducibility archive preserves the six final SNP2Cluster-Py v1.1.2 analyses, including the effective YAML configurations, redistributable analytical inputs, machine-readable results, validation and audit records, run manifests, static figures, interactive HTML visualizations and scientific HTML reports.

## Supplementary figures

- Figure S1: Complete Klebsiella pneumoniae annotated Core SNP heatmap from the full 172-isolate ST-only Core analysis.
- Figure S2: Complete Klebsiella pneumoniae annotated Core SNP heatmap from the full 172-isolate Facility + ST Core analysis.
- Figure S3: Facility-level Klebsiella pneumoniae transmission scatterplots for Hospitals A, B, D, E and F. The corresponding Hospital C analysis is presented in main-text Figure 3.
- Figure S4: Complete Streptococcus pneumoniae annotated Core SNP heatmap from the ST-only Core analysis.
- Figure S5: Complete Streptococcus pneumoniae annotated Core SNP heatmap with Community strain-sharing cluster annotations.

## Supplementary tables

Tables S1–S5 are provided as separate worksheets in SNP2Cluster-Py_Supplementary_Tables.xlsx.

- Table S1: Input datasets, effective analytical configurations, software version and run provenance for the six final analyses.
- Table S2: Context-level authoritative Core SNP-clustering results.
- Table S3: Facility-level Klebsiella pneumoniae Core and transmission-cluster summaries.
- Table S4: Community-level pneumococcal Core and strain-sharing cluster summaries.
- Table S5: Isolate-level comparison of ST-only and Facility + ST Klebsiella pneumoniae Core-cluster assignments.

## Supplementary interactive visualizations

Interactive Core minimum spanning trees and downstream Facility and Community networks are provided in their native HTML format in SNP2Cluster-Py_Interactive_Visualizations.zip. The archive preserves the directory structure of the corresponding final analysis runs and includes the machine-readable node, edge and cluster-assignment tables required to reconstruct the visualizations.

Core minimum spanning trees summarize genetic relationships among eligible isolates. Downstream network edges represent within-cluster minimum-spanning-tree relationships used for visualization and do not establish direct transmission, transmission direction or source attribution.

## Reproducibility archive

Complete reproducibility materials are available from the SNP2Cluster-Py manuscript companion repository. The repository contains the six final YAML configurations, redistributable analytical inputs, de-identified metadata, Core and downstream cluster tables, exploratory K-means and silhouette diagnostics, network node and edge tables, static figures, interactive HTML outputs, scientific HTML reports, figure-generation scripts and complete run instructions.

Each main-text and supplementary figure is linked to its corresponding analytical run, source files and generation script. Materials that cannot be redistributed are documented through data dictionaries, access conditions and analysis-reconstruction instructions. A versioned archive of the final reproducibility repository will be deposited in Zenodo, and the DOI will be added before submission.

