## Supplementary Figures for "SNP2Cluster-Py: A pathogen-agnostic, context-aware framework for Core SNP and transmission-cluster analysis"

### Figure S1

Complete Klebsiella pneumoniae annotated Core SNP heatmap from the full 172-isolate ST-only Core analysis.

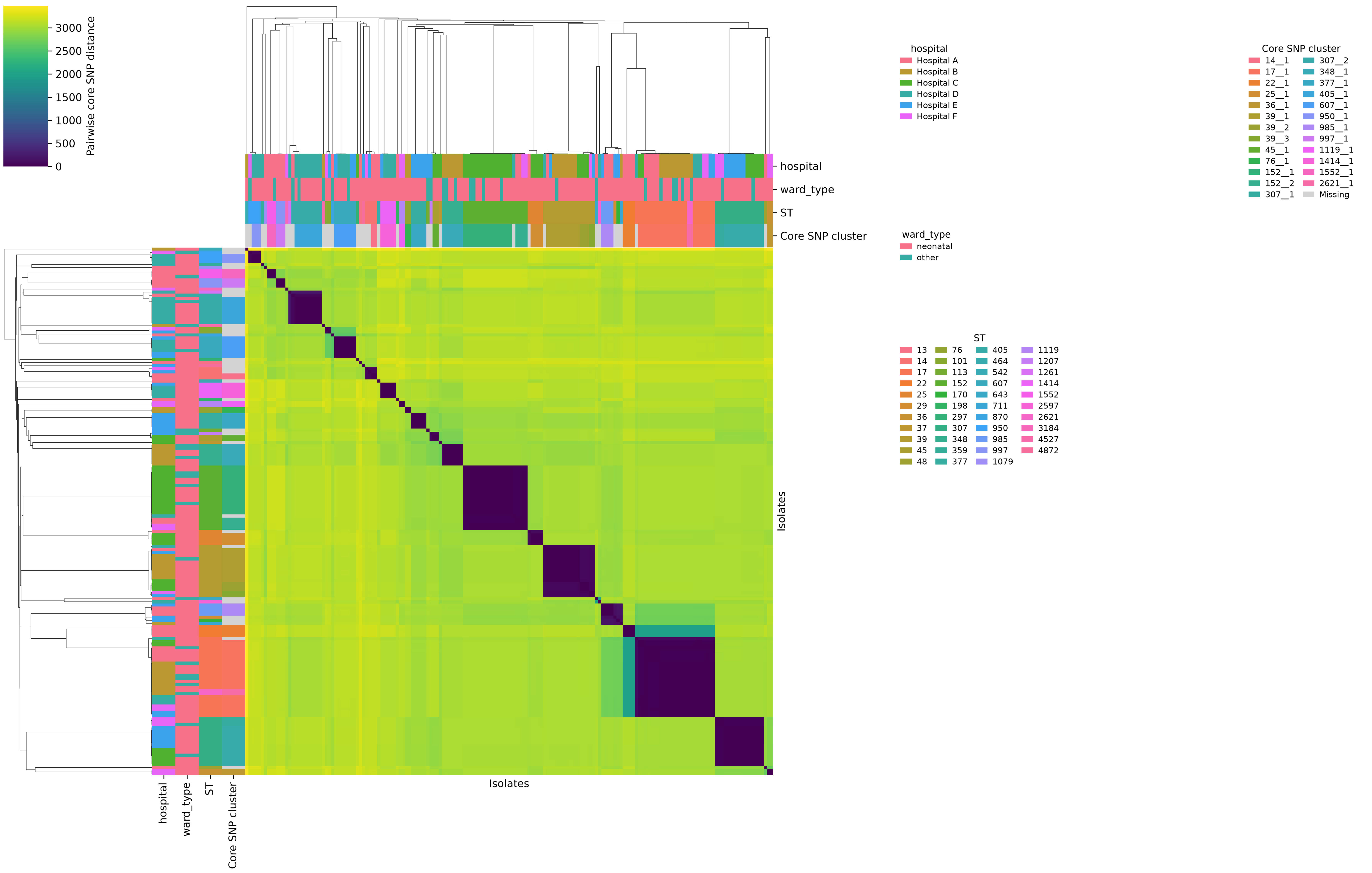

### Figure S2

Complete Klebsiella pneumoniae annotated Core SNP heatmap from the full 172-isolate Facility + ST Core analysis.

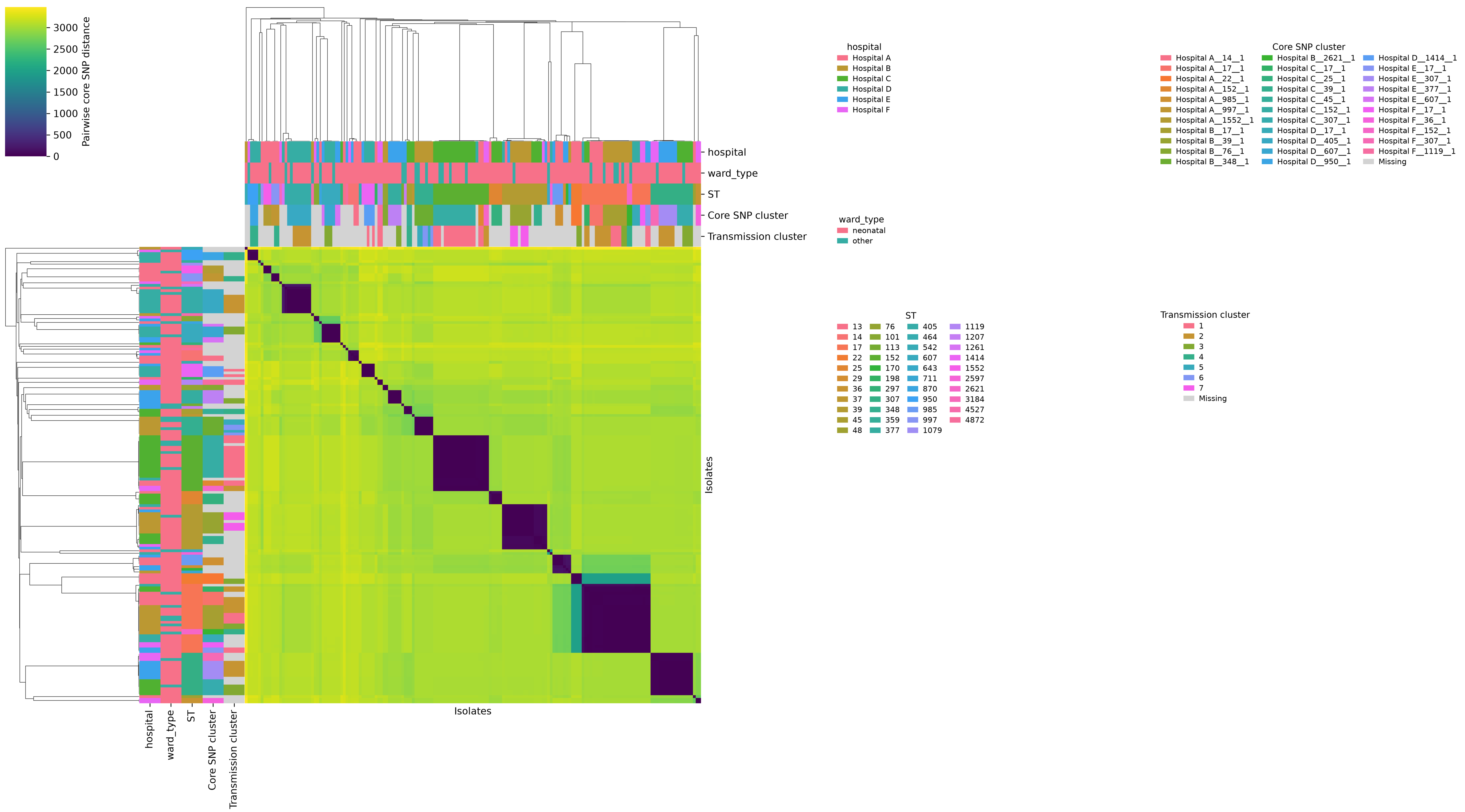

### Figure S3

Facility-level Klebsiella pneumoniae transmission scatterplots for Hospitals A, B, D, E and F. The corresponding Hospital C analysis is presented in main-text Figure 3.

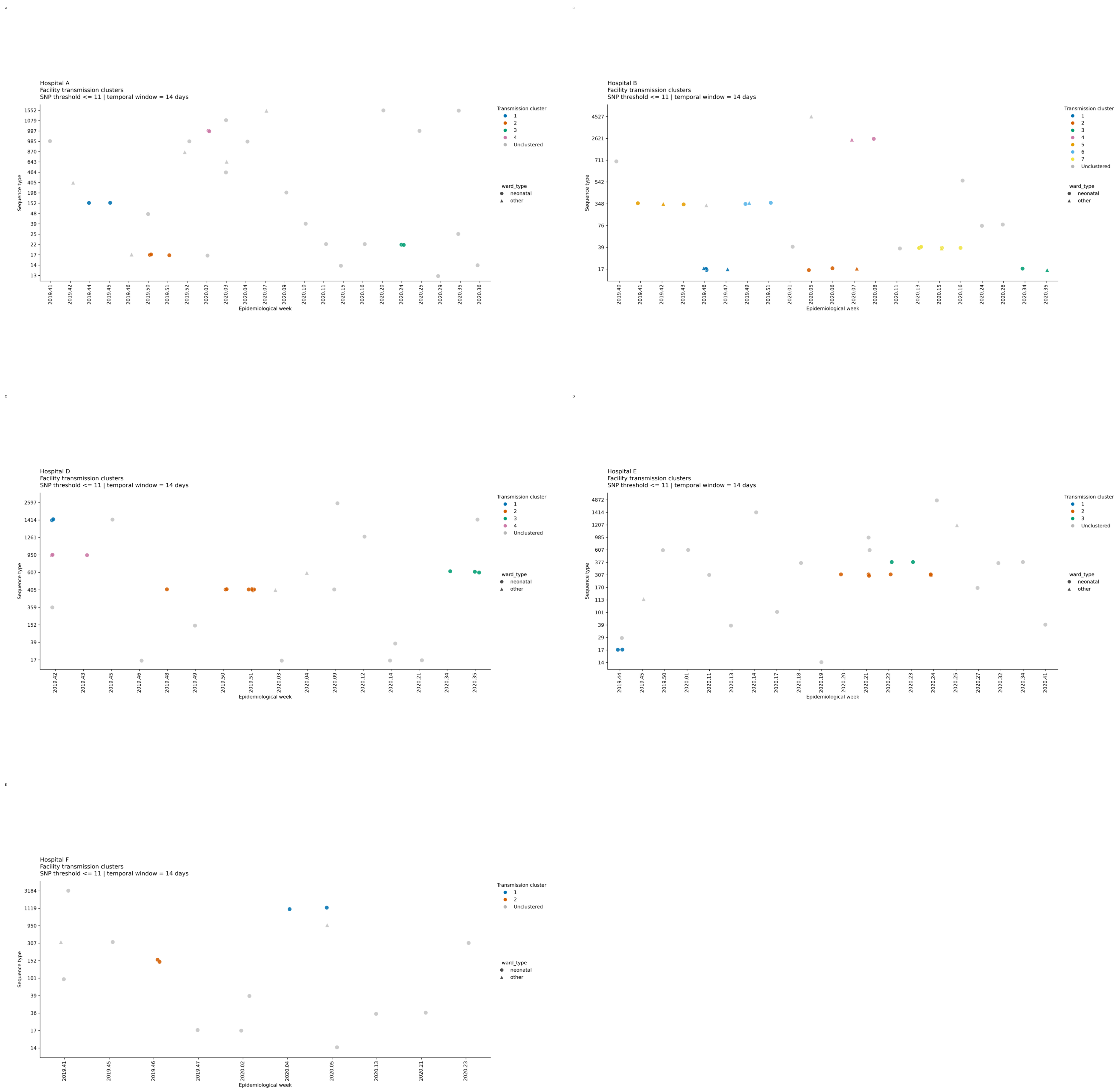

#### Figure S4

Complete Streptococcus pneumoniae annotated Core SNP heatmap from the ST-only Core analysis.

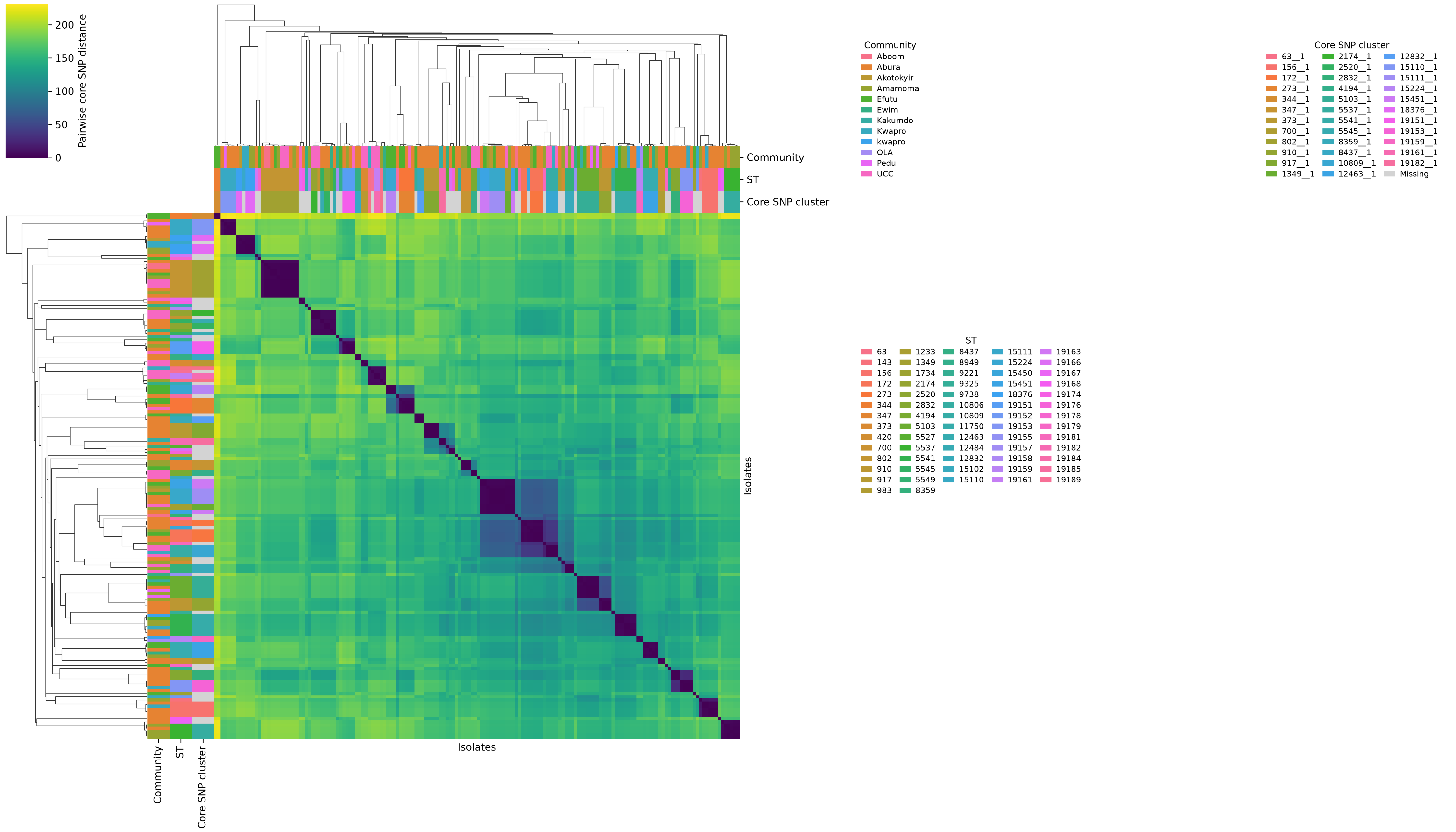

**Figure S5**

Complete Streptococcus pneumoniae annotated Core SNP heatmap with Community strain-sharing cluster annotations.

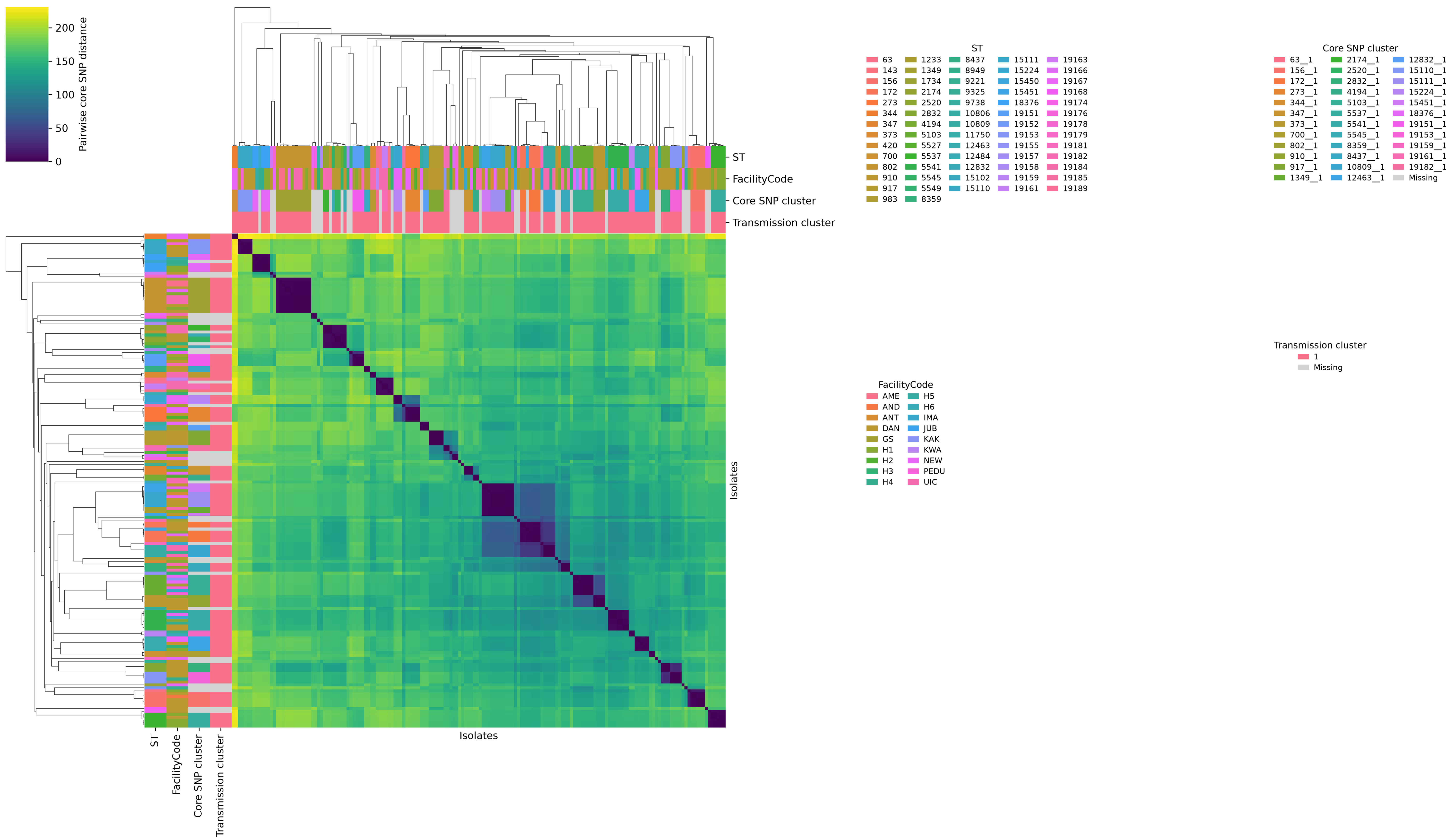
