## Supplementary HTML Files for "SNP2Cluster-Py: A pathogen-agnostic, context-aware framework for Core SNP and transmission-cluster analysis": 04_mst_Hospital_A.html

### Core SNP minimum spanning tree

hospital:
**Hospital A**

Core SNP cluster

14\_\_1

17\_\_1

22\_\_1

39\_\_1

152\_\_2

985\_\_1

997\_\_1

1552\_\_1

Missing

st\_type

13

14

17

22

25

39

48

152

198

405

464

643

870

985

997

1079

1552

Categories are encoded by the combination of
node shape and border pattern. Node fill colour
represents Core SNP cluster membership.

Lines represent minimum-spanning-tree connections.
Hover over an edge to view the SNP distance.
Hover over a node to view isolate metadata.
