## Supplementary HTML Files for "SNP2Cluster-Py: A pathogen-agnostic, context-aware framework for Core SNP and transmission-cluster analysis": 04_mst_Hospital_D.html

### Core SNP minimum spanning tree

hospital:
**Hospital D**

Core SNP cluster

17\_\_1

405\_\_1

607\_\_1

950\_\_1

1414\_\_1

Missing

st\_type

17

39

152

359

405

607

950

1261

1414

2597

Categories are encoded by the combination of
node shape and border pattern. Node fill colour
represents Core SNP cluster membership.

Lines represent minimum-spanning-tree connections.
Hover over an edge to view the SNP distance.
Hover over a node to view isolate metadata.
