## Supplementary HTML Files for "SNP2Cluster-Py: A pathogen-agnostic, context-aware framework for Core SNP and transmission-cluster analysis": 04_mst_Hospital_C.html

### Core SNP minimum spanning tree

hospital:
**Hospital C**

Core SNP cluster

Hospital C\_\_17\_\_1

Hospital C\_\_25\_\_1

Hospital C\_\_39\_\_1

Hospital C\_\_45\_\_1

Hospital C\_\_152\_\_1

Hospital C\_\_307\_\_1

Missing

st\_type

17

25

37

39

45

152

297

307

Categories are encoded by the combination of
node shape and border pattern. Node fill colour
represents Core SNP cluster membership.

Lines represent minimum-spanning-tree connections.
Hover over an edge to view the SNP distance.
Hover over a node to view isolate metadata.
