## Supplementary HTML Files for "SNP2Cluster-Py: A pathogen-agnostic, context-aware framework for Core SNP and transmission-cluster analysis": 05_transmission_network_Hospital_A.html

### Transmission-cluster network

hospital:
**Hospital A**

9 clustered isolates · 4 transmission clusters

Transmission cluster

1

2

3

4

ward\_type

neonatal

**How to interpret this network**  
  
Nodes represent isolates assigned to the same inferred transmission
cluster. Edges form a minimum spanning tree based on SNP distance
for visualization and do not represent direct transmission or
necessarily satisfy the configured temporal cutoff of
14 days.  
  
Transmission-cluster membership may reflect a qualifying chain
through intermediate isolates. Solid edges connect isolates whose
direct collection-date difference is within the temporal cutoff.
Dashed edges connect isolates whose direct collection-date
difference exceeds the cutoff.  
  
Review the corresponding transmission scatterplot for the temporal
distribution and possible time-based chain of cases. Hover over
nodes and edges for metadata and distance details.
