## Supplementary HTML Files for "SNP2Cluster-Py: A pathogen-agnostic, context-aware framework for Core SNP and transmission-cluster analysis": 04_mst_Abura.html

### Core SNP minimum spanning tree

Community:
**Abura**

Core SNP cluster

156\_\_1

172\_\_1

273\_\_1

700\_\_1

802\_\_1

910\_\_1

917\_\_1

2520\_\_1

2832\_\_1

5103\_\_1

5537\_\_1

5541\_\_1

5545\_\_1

8437\_\_1

12463\_\_1

12832\_\_1

15110\_\_1

15111\_\_1

15451\_\_1

18376\_\_1

19151\_\_1

19153\_\_1

19182\_\_1

Missing

ST

156

172

273

700

802

910

917

983

1233

2520

2832

5103

5537

5541

5545

8437

9221

9325

10806

12463

12832

15110

15111

15450

15451

18376

19151

19153

19163

19166

19167

19168

19174

19182

19184

19185

Categories are encoded by the combination of
node shape and border pattern. Node fill colour
represents Core SNP cluster membership.

Lines represent minimum-spanning-tree connections.
Hover over an edge to view the SNP distance.
Hover over a node to view isolate metadata.
