## Supplementary HTML Files for "SNP2Cluster-Py: A pathogen-agnostic, context-aware framework for Core SNP and transmission-cluster analysis": 04_mst_Amamoma.html

### Core SNP minimum spanning tree

Community:
**Amamoma**

Core SNP cluster

63\_\_1

172\_\_1

347\_\_1

802\_\_1

910\_\_1

1349\_\_1

5103\_\_1

5537\_\_1

5541\_\_1

8359\_\_1

12463\_\_1

15451\_\_1

18376\_\_1

19151\_\_1

Missing

ST

63

143

172

347

420

802

910

1349

5103

5537

5541

8359

12463

15451

18376

19151

19158

19181

Categories are encoded by the combination of
node shape and border pattern. Node fill colour
represents Core SNP cluster membership.

Lines represent minimum-spanning-tree connections.
Hover over an edge to view the SNP distance.
Hover over a node to view isolate metadata.
