## Supplementary HTML Files for "SNP2Cluster-Py: A pathogen-agnostic, context-aware framework for Core SNP and transmission-cluster analysis": 04_mst_Efutu.html

### Core SNP minimum spanning tree

Community:
**Efutu**

Core SNP cluster

172\_\_1

273\_\_1

344\_\_1

347\_\_1

802\_\_1

5103\_\_1

5541\_\_1

12463\_\_1

15111\_\_1

15224\_\_1

15451\_\_1

Missing

ST

172

273

344

347

802

1734

5103

5541

8949

9738

12463

15111

15224

15451

19176

19178

19179

Categories are encoded by the combination of
node shape and border pattern. Node fill colour
represents Core SNP cluster membership.

Lines represent minimum-spanning-tree connections.
Hover over an edge to view the SNP distance.
Hover over a node to view isolate metadata.
