## Supplementary HTML Files for "SNP2Cluster-Py: A pathogen-agnostic, context-aware framework for Core SNP and transmission-cluster analysis": 04_mst_Ewim.html

### Core SNP minimum spanning tree

Community:
**Ewim**

Core SNP cluster

12463\_\_1

Missing

ST

5103

5527

11750

12463

12484

19155

19157

Categories are encoded by the combination of
node shape and border pattern. Node fill colour
represents Core SNP cluster membership.

Lines represent minimum-spanning-tree connections.
Hover over an edge to view the SNP distance.
Hover over a node to view isolate metadata.
