## Supplementary HTML Files for "SNP2Cluster-Py: A pathogen-agnostic, context-aware framework for Core SNP and transmission-cluster analysis": 04_mst_UCC.html

### Core SNP minimum spanning tree

Community:
**UCC**

Core SNP cluster

273\_\_1

373\_\_1

802\_\_1

1349\_\_1

2174\_\_1

4194\_\_1

8359\_\_1

10809\_\_1

19151\_\_1

19161\_\_1

Missing

ST

273

373

802

1349

2174

4194

8359

10809

15102

19151

19161

19167

19189

Categories are encoded by the combination of
node shape and border pattern. Node fill colour
represents Core SNP cluster membership.

Lines represent minimum-spanning-tree connections.
Hover over an edge to view the SNP distance.
Hover over a node to view isolate metadata.
