## Supplementary HTML Files for "SNP2Cluster-Py: A pathogen-agnostic, context-aware framework for Core SNP and transmission-cluster analysis": 04_mst_5103.html

### Core SNP minimum spanning tree

ST:
**5103**

Core SNP cluster

5103\_\_1

Missing

FacilityCode

GS

H3

IMA

KAK

NEW

PEDU

Categories are encoded by the combination of
node shape and border pattern. Node fill colour
represents Core SNP cluster membership.

Lines represent minimum-spanning-tree connections.
Hover over an edge to view the SNP distance.
Hover over a node to view isolate metadata.
