## Supplementary HTML Files for "SNP2Cluster-Py: A pathogen-agnostic, context-aware framework for Core SNP and transmission-cluster analysis": 04_mst_19153.html

### Core SNP minimum spanning tree

ST:
**19153**

Core SNP cluster

19153\_\_1

FacilityCode

DAN

H4

Categories are encoded by the combination of
node shape and border pattern. Node fill colour
represents Core SNP cluster membership.

Lines represent minimum-spanning-tree connections.
Hover over an edge to view the SNP distance.
Hover over a node to view isolate metadata.
